# Polyamine elevation and nitrogen stress are toxic hallmarks of chronic sleep loss in *Drosophila melanogaster*

**DOI:** 10.1101/2021.10.01.462746

**Authors:** Joseph Bedont, Anna Kolesnik, Dania Malik, Aalim Weljie, Amita Sehgal

## Abstract

Chronic sleep loss profoundly impacts health in ways coupled to metabolism; however, much existing literature links sleep and metabolism only on acute timescales. To explore the impact of chronically reduced sleep, we conducted unbiased metabolomics on heads from three *Drosophila* short-sleeping mutants. Common features included elevated ornithine and polyamines; and lipid, acyl-carnitine, and TCA cycle changes suggesting mitochondrial dysfunction. Biochemical studies of overall, circulating, and excreted nitrogen in sleep mutants demonstrate a specific defect in eliminating nitrogen, suggesting that elevated polyamines may function as a nitrogen sink. Both supplementing polyamines and inhibiting their synthesis with RNAi regulated sleep in control flies. Finally, both polyamine-supplemented food and high-protein feeding were highly toxic to sleep mutants, suggesting their altered nitrogen metabolism is maladaptive. Together, our results suggest polyamine accumulation specifically, and nitrogen stress in general, as potential mechanisms linking chronic sleep loss to adverse health outcomes.

## Introduction

Sleep is a near-ubiquitous behavior in the animal kingdom, from humans on down to organisms as simple as hydra ^1^. The importance of sleep is evident in the consequences of neglecting this fundamental need. Chronic sleep deprivation (SD) is associated with a wide range of health maladies, up to and including death ^2^. Many theories of why sleep is so critical for health involve its links to metabolism. Some of these links are obvious; for instance, sleep has been proposed to allow recovery of energy stores and repair/replacement of macromolecules damaged during waking, both of which require dietary building blocks to happen ^2^. Other functions of sleep, like its proposed role in regulating synaptic strength and memory, are less obviously linked to metabolism; however, the potentiating effects of sleep on memory depend upon feeding status ^3,4^.

Importantly, while only chronic SD has been clearly coupled to most pathological outcomes of sleep loss, to date most metabolomic data available in chronic SD is from human insomnia or obstructive sleep apnea datasets, in which confounding variables are common and options for harvesting tissue are limited ^2,5^. Conversely, most animal model and all healthy human studies in this vein have examined the consequences of relatively acute, but not chronic, SD on the metabolome of various tissues and bodily fluids ^5^. This prompted us to ask the question: how does chronic loss of sleep affect metabolism, especially pathways relevant to pathologies such as neurodegeneration or behavioral impairment?

The fruit fly *Drosophila melanogaster* has been invaluable for determining the neurotransmitters and other intercellular signaling mechanisms that regulate sleep ^6^. Several sleep mutants with dramatically different mechanisms of action emerged from unbiased genetic screens, including *fumin* (*fmn*: defective dopamine reuptake), *redeye* (*rye*: nicotinic acetylcholine receptor loss-of-function), and *sleepless* (*sss*: dysregulated membrane potential and excessive GABA degradation) ^7–10^. Importantly, these mechanisms are well-conserved regulators of sleep-wake in mammals ^11^. In addition to pinpointing mechanisms regulating sleep amount, *Drosophila* has been an increasingly fruitful model for discerning how sleep regulates aspects of metabolism, including lipid homeostasis and autophagy, that are highly conserved across vertebrates ^12,13^.

The availability of the sleep mutants mentioned above, with profound sleep loss resulting from different mechanisms, provides an opportunity to uncover common features of chronic sleep loss. We conducted a metabolomic screen and report here that nitrogen metabolism is prominent among metabolites impaired across the mutants. In particular, sleep mutants have elevated ornithine and polyamines which arise, at least in part, from defective nitrogen excretion. Manipulating polyamines affects sleep amount, perhaps reflecting compensatory mechanisms regulating sleep in these mutants. But overall, the mutants’ remodeled nitrogen processing constitutes a pathological state that renders them very sensitive to dietary nitrogen. We suggest that polyamines specifically, and nitrogen metabolism in general, couple chronic SD to pathological outcomes like neurodegeneration and reduced lifespan. The link we demonstrate between polyamines and sleep also provides a mechanism for the integration of food and sleep homeostasis at the cellular level.

## Methods

### Fly husbandry

Heavily used fly lines in the manuscript include *iso31* control; sleep mutants *fmn, rye*, and *sss* on an *iso31* background (minimum 5X generations), and geneswitch/dicer lines were well-established in the lab prior to this study. RNAi alleles were ordered from Bloomington Drosophila Stock Center in Indiana, Vienna Drosophila Resource Center in Austria, or Kyoto Stock Center in Japan. See Table S2 for RNAi details of each RNAi line used in the manuscript.

### Metabolon global metabolomics and lipidomics

Ten total pools of ∼200-250 heads from mixed sex flies aged ∼1-2 weeks post-eclosion were collected for each genotype, split evenly between HD4 global metabolomics and CLP lipidomics assays. Collections were done at ∼ZT6; mid-day timepoint was chosen to enrich for metabolites dysregulated by chronic, as opposed to acute, sleep loss. Heads were collected by vortexing whole flies snap frozen on dry ice and separating heads from bodies by size on dry ice-cooled grates. Samples were stored at -80C and shipped to Metabolon on dry ice. Sample preparation, control procedures, and analysis were carried out at Metabolon Inc as described elsewhere ^14–18^. Both HD4 and CLP procedures are briefly outlined below.

### HD4 Global Metabolomics

Samples extracted and spiked with recovery standards using a MicroSTAR System (Hamilton Company) and methanol-precipitated under vigorous shaking Genogrinder 2000 (Glen Mills). Samples were fractionated, dried, resuspended in appropriate solvents, and analyzed using four distinct modes on a Waters ACQUITY ultra-performance liquid chromatography (UPLC) and a Thermo Scientific Q-Exactive high resolution/accurate mass spectrometer interfaced with a heated electrospray ionization (HESI-II) source and Orbitrap mass analyzer operated at 35,000 mass resolution. Metabolites were identified by the Laboratory Information Management System, an automated system that identified ion features in our head lysate samples using a reference library of known metabolites defined by retention time, molecular weight (m/z), preferred adducts, in-source fragments, and associated MS spectra. The data was curated by visual quality control using software developed at Metabolon.

### CLP Lipidomics

Lipids were extracted using a modified Bligh-Dyer extraction method with deuterated internal standards. Samples were then subjected to infusion-MS analysis in both positive and negative modes on a Shimadzu LC with nano PEEK tubing and a Sciex SelexIon-5500 QTRAP in MRM mode (>1,100 MRMs). Individual lipids were quantified as signal / internal standard and summed into class and total lipid concentrations.

### Acute SD targeted nitrogen metabolomics

Twenty total pools of ∼90 mixed sex *iso31* flies aged ∼1.5 weeks post-eclosion were divided evenly among four conditions: (1) collected at ZT2, (2) collected at ZT14, (3) collected at ZT2 after a 14hr mechanical sleep deprivation, and (4) collected at ZT2 after a 12hr sleep deprivation followed by a 2hr sleep rebound. Heads were collected by vortexing whole flies snap frozen on dry ice and separating heads from bodies by size on dry ice-cooled grates. Samples were stored at -80C until they were processed. Samples were extracted and prepared for LC-MS analysis as previously described ^19,20^. Briefly, a stainless steel bead and 300 μl of 2:1 Methanol:Chloroform were added to each sample. Samples were homogenized for a total of 4 minutes at 25 Hz in a tissue homogenizer. Next, 100 μl of water and chloroform were added to each sample. Samples were vortexed and then centrifuged for 10 minutes at 13,300 rpm at 4°C. 170 μl of the upper fraction containing polar metabolites was collected from each sample and dried in a speed vacuum for 2.5 hours. Dried fractions were resuspended in 100 μl of acetonitrile:water, vortexed for 20 seconds and centrifuged for 10 minutes at 13,300 rpm at 4°C prior to transferring to MS vials. Samples were analyzed in analytical triplicates and pooled quality control samples were run at the beginning and end of the run as well as after every 6th injection. For each sample, 2 μl were injected onto an Acquity UPLC BEH Amide column (1.7 μm, 2.1 mm × 150 mm) with a 0.2 μm inline precolumn filter using an Acquity H-Class UPLC system (Waters Corporation) coupled to a Xevo TQ-S micro mass spectrometer operating in a positive ion polarity mode. Initial chromatographic conditions consisted of 100% Solvent D (90:10 Acetonitrile:water, 2 mM Ammonium Acetate, 0.2% Formic Acid) ramped to 79.4% Solvent A (95:5 milliQH2O:Acetontrile, 2 mM Ammonium Acetate, 0.2% Formic Acid) in 15 minutes. The column was washed in 100% Solvent A for 5.5 minutes before reequilibrating in 100% D. A total of 18 compounds were measured through targeted MRM methods. Transitions for Urea, Sarcosine, Proline, Trans-4-Hydroxyproline, Ornithine, Spermidine, Acetylornithine, Citrulline, SAM, Spermine, Glutamate, 4-Guanidinobutanoic Acid, Creatine, Acetylputrescine, and GABA were used as described in ^20^. Additional transitions were added for Argininosuccinate (291.13/70.07 (15/40) [(Cone Voltage/Collision Energy]), Arginine (157.14/60.05 (30/12)), and Putrescine (88.9/54.86 (14/16) and 89.11/72.08 (15/20)). Data was processed using TargetLynx (Waters) to obtain ion counts for further analysis using an in-house R-script. Spermine is excluded from our results because of low signal / high noise that rendered the signal suspect. Urea is excluded from our results because we were unable to corroborate its relevance through biochemical methods.

### Lysate collection from whole flies for biochemistry

Pools of 5 female or 5 male ∼1.5 weeks post-eclosion flies were anesthetized with CO2 and quickly sorted into 2mL Safelock Tubes, weighed, then snap frozen on dry ice. Blocks were pre-chilled to 4C, and the tubes were transferred to wet ice, where 200uL of 1X PBS supplemented with 1 cOmplete EDTA-free protease inhibitor tablet / 2.5mLs PBS (PBS-PI) and a stainless steel bead were added. Samples were quickly loaded into chilled blocks and lysed on a TissueLyser II (Qiagen) at 25.0m/sec in two 2-min bursts. Beads were removed and solid detritus was pelleted by spinning at 15,000rpm for 15min at 4C. Supernatant was carefully transferred to a clean tube and diluted 1:4 in PBS-PI before being used for total protein, ammonia, and urate (uric acid) biochemical assays.

### Hemolymph collection from female flies for biochemistry

Only female flies were run for these experiments because their typical hemolymph volume is much higher than males. Groups of 25 ∼1.5 weeks post-eclosion flies were anesthetized with CO_2_, rapidly pricked in the thorax with tungsten probes (Ted Pella 13570), and loaded into 0.5mL Ependorfs perforated at the base with 22-gauge syringe tips, which were nested inside of 1.5mL Ependorfs containing 45uL of PBS-PI. Nested tubes were then centrifuged at 5000rpm for 5min at 4C. 25uL of hemolymph + PBS-PI was removed for total protein analysis, and the remaining ∼20uL was diluted with a further 60uL of PBS-PI for ammonia analysis. Flies were weighed after hemolymph harvest.

### Excrement collection from male flies for biochemistry

Only male flies were run for these experiments to eliminate egg-laying as a confound. Groups of 12 ∼1.5 weeks post-eclosion flies were anesthetized with CO2 and sorted into 1.5mL Ependorf tubes perforated twice through each cap with an 18-gauge needle. The flies were returned to their home incubators for 2 hours, then anesthetized with CO2 and flipped into fresh 1.5mL Ependorf tubes to be weighed. The excrement in the first set of Ependorfs was resuspended by vortexing into 150uL of PBS-PI, which was used at this concentration to run ammonia and urate (uric acid) biochemical assays.

### Biochemical Assays: Total Protein, Ammonia, Uric Acid, Urea

Total protein assay was conducted using an Abcam 207003 total protein assay kit according to manufacturer instructions: absorbance measured at 540nm. Ammonia assay was conducted using a Sigma-Aldrich MAK310 kit according to manufacturer instructions: fluorescence measured at excitation 355nm / emission 460nm. Uric acid assay was conducted using a Sigma-Aldrich MAK077 kit according to manufacturer instructions: fluorescence measured at excitation 535nm / emission 595nm. Signal from all biochemical assays was normalized to body weight of the pools of flies that provided the material. Fluorescence and absorbance were measured using a Victor-3V plate reader (Perkin-Elmer) or a Cytation 5 (BioTek).

### Blue-Poo Assay

Male flies were pre-fed on our lab’s standard yeast-molasses food supplemented with 2.5mg/mL of FD&C blue 1 / FCF brilliant blue dye for 24 hours. Excrement was then collected as though for biochemistry, then resuspended into 150uL of 1X PBS. Absorbance was measured at 620nm and normalized to body weight to calculate fecal volume.

### Sleep Experiments

For polyamine supplementation experiments, ∼3-5 days post-eclosion flies of both sexes were loaded into locomotor tubes with 5% sucrose / 2% agar food supplemented with water vehicle, 16mM L-ornithine monohydrochloride (Sigma-Aldrich 2375), 16mM putrescine dihydrochloride (Sigma-Aldrich P7505), or 16mM spermidine trihydrochloride (Sigma-Aldrich S2501). Sleep was measured from movements on DAM5H multibeam monitors (Trikinetics), averaged across the 2^nd^-4^th^ full days of recording. Four independent experiments were pooled to generate the data shown.

For nitrogen metabolism RNAi screen, ∼3-5 days post-eclosion female flies were loaded into locomotor tubes with 5% sucrose / 2% agar food supplemented with 500uM mifepristone (RU+ food) (Sigma-Aldrich M8046). Sleep was recorded from counts of beam breaks on single-beam DAM2 monitors (Trikinetics), and the 4^th^-5^th^ full days of exposure to RU+ food were averaged to determine sleep. Geneswitch(GS)>Dicer, RNAi crosses with mean sleep at least 60 min higher or lower than both GS>Dicer and RNAi controls were considered potential hits and validated. Validation of promising crosses was carried out similarly to the screen, but included both RU+ and ethanol vehicle (RU-) food conditions to assess whether effects were acute. If the first validation experiment was inconsistent with the initial screen result for a given RNAi, further validation of that RNAi was terminated. At least two independent validation experiments were run for each nitrogen RNAi cross identified as having a legitimate sleep phenotype.

DAMfilescan and custom MatLab scripts were used to calculate sleep metrics for both sets of experiments ^21^.

### qPCR Validation of Nitrogen Pathway RNAis

To confirm knockdown of target transcripts with our nitrogen metabolism RNAis that yielded sleep phenotypes, we crossed these RNAi’s together with uas-dicer/+;actinGS/+ and closely followed the same whole-fly RNA collection, cDNA synthesis, and qPCR method we published previously ^13^. The following qPCR primer sets were used for each target transcript: *asl* forward: TCGACAAGCTGTCCCAAGTG reverse: CACCAGATAGTAGGCCCAGTC *spds* forward: GAAACACGCGCTGAAGGATG reverse: GGCATAGGCCACCTTAGCAA *sms* forward: GAGCTGCAGAACATTGCTGA reverse: GTACAACAAGGCGCCATCAC *odc2* forward: CTATGCCGTCAAGTGCAACG reverse: CCAAGTCCCAGGACCAACTT *odc1* forward: CCCAACTCCAACCTGATCGT reverse: AGTAGTGCGCCGAACTTGAA *α-tubulin* forward: CGTCTGGACCACAAGTTCGA reverse: CCTCCATACCCTCACCAACGT

### Lifespan Experiments

For all lifespan studies, mated flies of both sexes that had eclosed within the preceding ∼2 days were collected under CO2 anesthesia and housed single-sex on standard or special food at a maximum density of 30/vial.

For classical lifespan studies, flies were flipped to fresh vials and dead were tallied every 3 days (experiment comparing genotype lifespan on standard food) or every 2 days (experiment comparing within-genotype lifespan on high protein, high sugar, and only sugar food). Each vial was longitudinally maintained on the same assigned food condition, and dead flies were counted at each flip, tracking survival in this way until all flies were dead. Flies were never again anesthetized after initial collection.

For starvation-challenge lifespan studies, flies were primed on high protein, high sugar, or only sugar diet for 5 days, flipping onto fresh food midway through the priming period. All flies were then anesthetized with CO2 and loaded into locomotor tubes containing 2% agar with no sugar. Behavior was recorded using single-beam DAM monitors until all flies were dead. Sleep records were analyzed using custom Matlab scripts (Hsu et al, 2020) to determine survival time of each fly with half-hour precision, defining death as the first half-hour bin of apparent ‘sleep’ that did not end. Flies that died during loading before the start of our sleep recording were excluded from the analysis.

For survival on nitrogenous metabolite supplementation studies, flies were initially maintained on standard food, then loaded into locomotor tubes containing 5% sucrose / 2% agar food drugged with either water vehicle, ornithine, or polyamine at ∼3-4 days post-eclosion. Behavior was recorded using single-beam DAM monitors until the end of the fifth complete day on supplemented food, plus the balance of the loading day. Sleep records were analyzed similarly to starvation-challenge, expect that all flies surviving past the fifth full day of recording were censored.

Sehgal Lab Standard Yeast-Molasses Diet: 64.7g/L corn meal ; 27.1g/L dry yeast ; 8g/L agar ; 61.6mL/L molasses ; 10.2mL/L 20% tegosept ; 2.5mL/L propionic acid

High-Protein Diet: Sehgal Lab Standard Yeast-Molasses + extra 116g/L yeast

High-Sugar Diet: Sehgal Lab Standard Yeast-Molasses + extra 116g/L sucrose

All-Sugar Diet: 27% w/v sucrose-agar.

### Blue Dye Feeding Assay

Flies were adapted to high-protein, high-sugar, or all-sugar diet for 5 days before measuring feeding. At ZT0 of the sixth day, flies were flipped onto the adapted diet laced with 2.5mg/mL FD&C blue 1 / FCF brilliant blue dye for 4 hours. Whole fly lysates were then extracted using our procedure for biochemistry above, but using 1X PBS instead of PBS-PI as the lysis buffer. Absorbance was measured at 620nm and normalized to body weight to calculate ingested food volume.

## Results

### Chronic, but not acute, sleep loss up-regulates polyamine levels in *Drosophila* heads

To assess metabolic consequences of chronic sleep loss, we conducted unbiased global and lipid metabolomics on pools of heads collected at mid-day from *iso31* control and sleep mutant *fmn, rye*, and *sss* flies. Our global screen identified 30 metabolites commonly regulated across all sleep mutants, which we attribute to chronic sleep loss (p<0.05 / q<0.05; Figure 1A).

**Figure 1:**
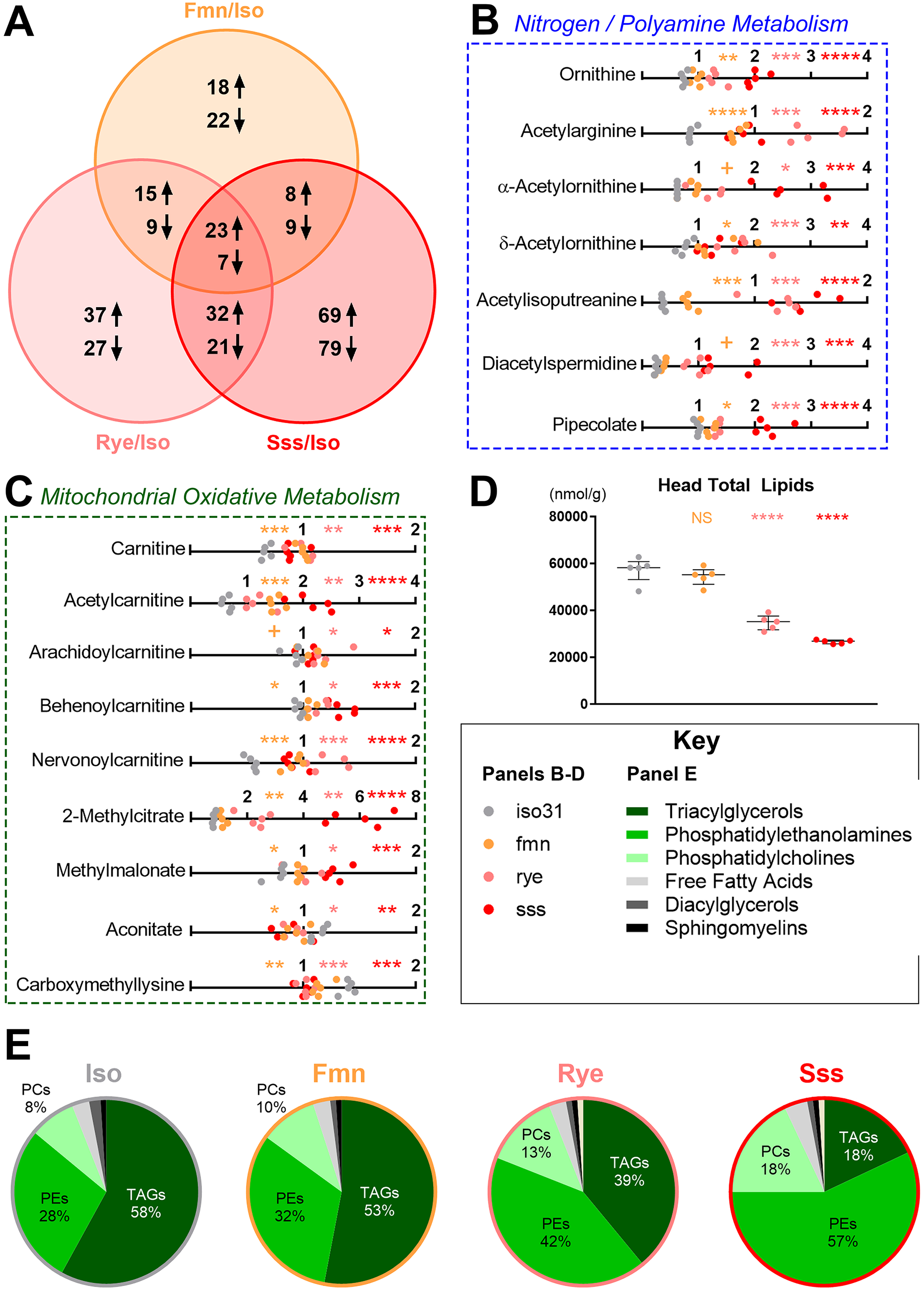
Sleep mutants have profoundly altered nitrogen, mitochondrial, and lipid metabolism. All metabolomic data is from *iso31* (gray), *fmn* (orange), *rye* (pink), and *sss* (red) pools of fly heads collected at ∼ZT6 on a 12hr:12hr light:dark cycle. All statistical comparisons shown are color-matched sleep mutant vs *iso31* control. (A) Venn diagram indicating the number of metabolites consistently either up- or down-regulated in one or more sleep mutants compared to *iso31* control. (B-C) Line graphs showing scaled metabolite levels in head lysates, grouped by involvement in nitrogen metabolism (B) or mitochondrial oxidative metabolism (C). Data shown are individual pools of fly head lysate; n=5; Welch’s t-tests (p-values) with FDR correction for multiple comparisons (q-values); + p<0.05 but q>0.05, * p/q<0.05, ** p/q<0.01, *** p/q<0.001, **** p/q<0.0001. (D) XY graph showing total lipid content in head lysates. Data shown are individual pools of fly head lysate overlaid with median+/-interquartiles; n=5; Dunnett test; NS = not significant, ****p<0.0001. (E) Pie graphs showing the percentage contributions of major lipid families to the head lipidome of each genotype. Changes in sleep mutant lipid composition are largely driven by decreased overall lipid content (D), which is disproportionately driven by decreased triacylglycerol levels, especially in *rye* and *sss*.

Of these, 7 metabolites (∼23%) are primarily linked to nitrogen detoxification and/or polyamine synthesis: ornithine, N-acetylarginine, N-α-acetylornithine, N-δ-acetylornithine, N-acetylisoputreanine, N1-N8-acetylspermidine, and pipecolate (q<0.05; Figure 1B). The major polyamines putrescine and spermidine were also significantly elevated in *rye* and *sss* (q<0.05), and putrescine is likely elevated in *fmn* as well (p=0.0535) (Figure S1A). Other commonly regulated metabolites, such as taurine, sarcosine, and guanine, have links to nitrogen metabolism, but also to other pathways (Figure S1B) ^22–24^. Overall, in sleep mutants, the nitrogen metabolome had selective up-regulation of ornithine, polyamines, and related species, with few consistent changes in metabolites in alternate pathways like urate synthesis, proline synthesis, or nitric oxide synthesis (Figure S1A; Table S1).

In a follow-up experiment, we asked if acute sleep deprivation alters nitrogen metabolism in a similar way. Targeted metabolomics examining levels of polyamines and directly linked pathways in *iso31* heads at ZT2 (control) compared to ZT14, ZT2 following overnight sleep deprivation (ZT2 SD), and ZT2 following overnight SD with an opportunity for morning rebound sleep (ZT2 SR), showed minimal effects on all nitrogen metabolites tested. Ornithine alone trended higher at ZT2 SR compared to ZT2 (p<0.05; q>0.05; Figure S1C). Thus, the remodeled nitrogen metabolome observed in sleep mutants is a specific consequence of chronic, not acute, sleep loss.

### Mitochondrial, lipid, and other metabolome changes in heads of *Drosophila* sleep mutants

Another 5 metabolites commonly regulated across sleep mutants (carnitine, acetyl-carnitine, arachidoylcarnitine, behenoylcarnitine, and nervonoylcarnitine) suggested probable defects in mitochondrial beta-oxidation (Figure 1C). Other acyl-carnitines were elevated idiosyncratically, with shorter-chained species accumulating in *fmn* and *rye*, while *sss* was depleted of these, instead accumulating longer-chained acyl-carnitines (Table S1). Elevated 2-Methylcitrate and methylmalonate, and lower N6-carboxymethyllysine and aconitate levels are consistent with disrupted oxidative mitochondrial energy production across sleep mutants (q<0.05; Figure 1C) ^25–28^. In *rye* and *sss*, mitochondrial defects likely contribute to overall lipid loss, and bias the *rye* and *sss* lipidome dramatically toward phosphatidylcholines and phosphatidylethanolamines; *fmn* has weak trends in a similar direction (Figure 1D-E). Cholesteryl esters as a class are also disrupted, with decreases in *rye* and *sss* (q<0.05) and a similar trend in *fmn* (p<0.05) (Table S1).

The remaining metabolites commonly regulated across sleep mutants were more varied, but included two smaller clusters of related metabolites: threonine derivatives (threonate, N-acetylthreonine, and gamma-glutamylthreonine) and erythronate/erythritol (Figure S1B).

### *Drosophila* sleep mutants have a common defect in nitrogen excretion

Metabolomes of sleep mutant heads had no consistently down-regulated pathways that corresponded to the elevation of ornithine and many polyamine species. Thus, we suspected that polyamine elevation serves as a sink for surfeit nitrogen waste such as urate, ammonia and urea. While urate was not consistently different in sleep mutant heads (Table S1), ammonia and urea were not detected by Metabolon, so we tested this biochemically. Unable to reliably detect nitrogen waste metabolites in heads biochemically (data not shown), we tested total protein, ammonia, urate, and urea levels in whole fly. Urea was unquantifiable over interference from other nitrogen metabolites even in whole-fly samples (data not shown), which is consistent with urate being the primary nitrogen waste product in terrestrial insects, and with *melanogaster*’s lack of an obvious *otc* homolog to complete the urea cycle ^22^. However, total protein, ammonia, and urate levels in whole flies were similar in controls and sleep mutants, with the exceptions of urate elevation in *sss* of both sexes, and decreased urate in *fmn* males (Figure S2A-F). Curious whether differences in nitrogen waste might be confined to the fly’s circulation, we assayed hemolymph total protein and ammonia levels in sleep mutant and control flies at dawn and dusk. Total protein and ammonia were significantly elevated only in *rye* hemolymph (Figure S2G-H).

We reasoned that polyamine up-regulation and genotype-specific nitrogen metabolome changes might be successful in dissipating nitrogen stress, which would account for their lack of accumulated nitrogen waste metabolites. However, deficits in nitrogen processing may be reflected in excretions in this case. Thus, we tested whether sleep mutant flies were deficient at physiological elimination of toxic nitrogen species that cause nitrogen stress, by measuring urate and ammonia in fly excrement. Because excretion is an active behavior, we assayed it at the beginning and end of waking (ZT0-2 and ZT10-12), and in the middle of the night (ZT17-19). Each sleep mutant excreted significantly less urate at ZT0-2 (*rye/sss*) or ZT10-12 (*fmn*) with trends lower at all timepoints tested, including late at night when sleep mutants are more likely to be awake and able to excrete than controls (p<0.05; Figure 2A). Sleep mutants also trend toward lower direct excretion of ammonia at all timepoints tested, including significantly lower excretion at ZT10-12, when *iso31* excretes more ammonia than at other times tested (p<0.05; Figure 2B). Ongoing experiments measuring blue dye excretion after 24 hours of dye feeding at these same circadian times suggest that the sleep mutants do not have a consistent deficit in volume of their excretions (data not shown). All sleep mutants we tested thus have a consistent, specific deficit in nitrogen metabolite excretion. This metabolic correlate is both consistent across sleep mutants, and able to explain elevated polyamines.

**Figure 2:**
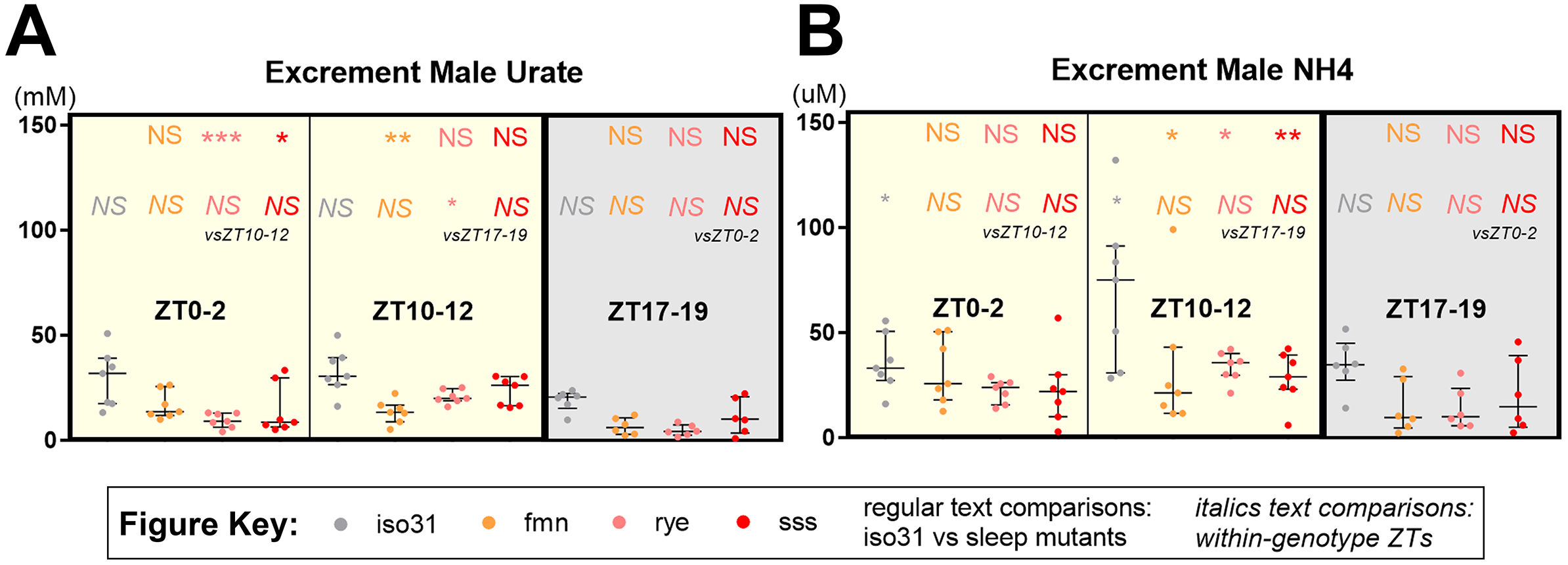
Sleep mutants are deficient at excreting nitrogen waste metabolites. All biochemical data are from *iso31* (gray), *fmn* (orange), *rye* (pink), and *sss* (red) flies on a 12hr:12hr light:dark cycle. All top-row, regular-text statistical comparisons shown are for color-matched sleep mutant vs iso31 control. All bottom-row, italicized statistical comparisons shown are for color-matched within-genotype time-of-day comparisons. (A-B) Excreted urate (A) and ammonia (B) at ZT0-2, ZT10-12, or ZT17-19. Data shown are level in excretions collected over the indicated 2-hour span from individual pools of male flies resuspended in PBS + protease inhibitor, overlaid with median+/-interquartiles; n=6-7. All experiments used Tukey HSD tests; NS = not significant, * p<0.05, ** p<0.01, *** p<0.001, **** p<0.0001.

### Blocking polyamine synthesis increases sleep in *Drosophila melanogaster*

Both feeding ornithine and blocking ornithine’s entry into proline synthesis by knocking down *oat* in neurons have been reported to increase sleep in Drosophila ^1^. Since sleep loss is thought to promote accumulation of somnogens, we wondered whether our metabolomic screen, together with these prior results, were hinting at a sleep-promoting role for polyamines, since their production could theoretically be enhanced by both *oat* knockdown and ornithine supplementation. We conducted RNAi screens targeting a range of enzymes involved in polyamine synthesis and linked pathways, including proline synthesis and nitric oxide synthesis, to broadly test how ornithine-based nitrogen metabolism regulates sleep in *Drosophila*. The use of drug-inducible geneswitch drivers allowed us to restrict manipulations to adulthood.

We first screened with actin-geneswitch (GS) uas-dicer on RU+ food, knocking down expression of the target genes in the whole fly and measuring sleep. This screen yielded single RNAi hits for three genes: the arginine synthesizing enzyme argininosuccinate lyase (*asl*), and the polyamine synthesis enzymes spermidine synthase (*spds*) and spermine synthase (*sms*) (Figure 3A). We followed up by screening the same pool of RNAis with nsybGS uas-dicer, knocking down the target genes pan-neuronally. This screen yielded only a single hit: an RNAi for ornithine decarboxylase 2 (*odc2*) (Figure 3B). Neither screen recapitulated the previous report of sleep gain with pan-neuronal elav-Gal4 knockdown of *oat*, with either the RNAi used in that study or two alternatives (Table S2) ^1^.

**Figure 3:**
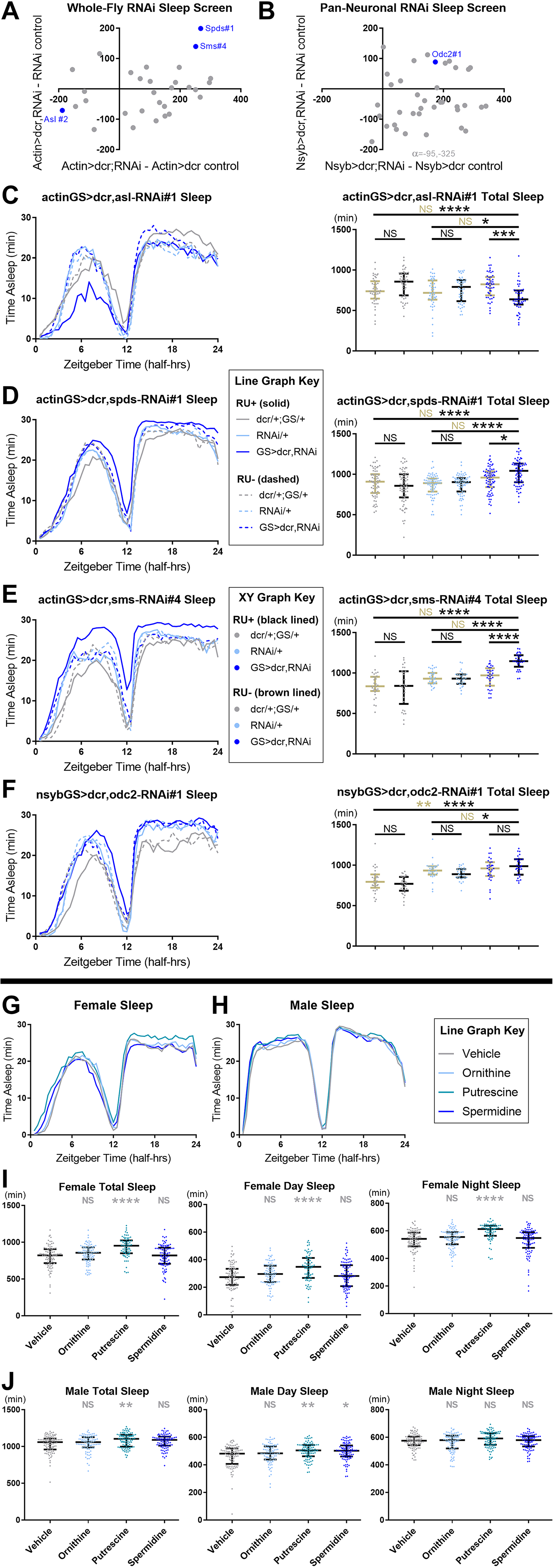
Sleep is increased by both supplementing and inhibiting synthesis of polyamines. All behavioral data was collected from flies on a 12hr:12hr light:dark cycle. (A-B) Difference in first-pass population mean sleep on mifepristone-laced (RU+) food for a range of female geneswitch (GS)> dcr;nitrogen metabolism-RNAi crosses compared with GS> dcr control (x-axis) and RNAi control (y-axis). Geneswitches are whole-fly actinGS in (A) and pan-neuronal nsybGS in (B). Blue, labeled dots indicate significant hits that passed all validation steps. RNAi identities and population statistics for the screens are provided in Table S2. (C-F) Sleep data from validation experiments of confirmed hits from the screens, shown as line graphs of averaged sleep behavior over circadian time (left) and statistical comparison of total sleep from individual flies overlaid with median+/-interquartiles (right). Whole-fly knockdown of argininosuccinate lyase (*asl*, C) RU-dependently decreased sleep. Whole-fly knockdown of spermidine synthase (*spds*, D) and spermine synthase (*sms*, E), as well as pan-neuronal knockdown of ornithine decarboxylase (*odc*, F) RU-dependently increased sleep. n=30-71 flies per group; Tukey HSD (E) or Steel-Dwass (C,D,F) tests; statistics for genotype comparisons are color-coded for vehicle control (RU-, brown) or RU+ (black). (G-H) Averaged sleep behavior of *iso31* mated females (G) or mated males (H) plotted over circadian time, on food supplemented with water vehicle (gray), 16mM L-ornithine (light blue), 16mM putrescine (medium blue), or 16mM spermidine (dark blue). (I-J) Total sleep (left), day sleep (middle), and night sleep (right) in female (I) and male (J) flies. Data shown is individual-fly averages overlaid with population median+/-interquartiles. The same dataset was used to compute G-J and auxiliary metrics shown in Table S2. n=69-93 flies per group; Steel test (vs Vehicle) used for most panels, Dunnett test (vs Vehicle) used for I-day. For all panels, NS = not significant, * p<0.05; ** p<0.01; ***p<0.001; **** p<0.0001.

Whole-fly *asl* RNAi decreased sleep (Figure 3C). This was driven primarily by decreased day sleep, though sleep was fragmented throughout the cycle; sleep latency was unaffected (Table S2). Whole-fly *spds* or *sms* RNAi robustly increased sleep throughout the cycle, with marked consolidation of sleep and decreased night sleep latency (Figure 3D-E; Table S2). Pan-neuronal *odc2* RNAi increased total and night sleep; there was neither significant consolidation nor decreased sleep latency associated with this sleep gain (Figure 3F; Table S2). None of these knockdowns produced changes in activity index that might confound their sleep phenotypes (Table S2). Finally, measuring whole-fly RNA expression for target transcripts with actinGS uas-dicer knockdown demonstrates that the sleep-affecting *asl, spds*, and *sms* RNAis robustly knock down their targets (Figure S3A-C). The sleep-promoting *odc2* RNAi does not measurably knock down *odc2*; however, it decreases expression of the higher-expressed homolog *odc1*, suggesting this knockdown mediates its sleep effects (Figure S3D-E).

Finding only single RNAis for particular genes in these pathways prevents us from excluding RNAi off-targets as an alternative explanation for any single gene’s effect on sleep. However, contrary to our hypothesis, sleep-promoting effects of single RNAis targeting three consecutive steps in the polyamine pathway suggest that blocking polyamine synthesis in general is sleep promoting.

### Feeding polyamines, but not ornithine, promotes sleep in control flies and is lethal to sleep mutants

Following up on our surprising RNAi sleep results, we tested the sleep effects of supplementing polyamines by assaying sleep in *iso31* control flies maintained on food supplemented with either water vehicle or 16mM L-ornithine, putrescine, or spermidine, tentatively hypothesizing that polyamines would be wake-promoting. The use of multibeam DAM5H monitors that provide high resolution analysis of locomotor activity allowed us to control for changes in fly locomotion or location that could be caused by toxicity or chemotactic properties of the metabolites ^29^.

Again contrary to our expectations, polyamines increased sleep in both mated female and male flies in our hands (Figure 3G-H). Specifically, putrescine increased total sleep and day sleep, and decreased latency to sleep during the day in both sexes; in addition, it tentatively increased night sleep and decreased latency to sleep at night in females (p<0.05; PA Figure 3I-J). Sleep gains with putrescine were driven primarily by consolidation of day sleep (p<0.0001; Table S2). Somewhat elevated attrition and a modest decrease in nighttime activity index in females on putrescine are caveats to whether they truly sleep more at night. However, putrescine neither increases attrition nor decreases activity index in males, nor does it decrease female daytime activity index, affirming daytime sleep gains (Table S2).

Spermidine supplementation had milder but similar effects on sleep. While insufficient to drive an increase in total sleep, spermidine increased day sleep in males, and decreased sleep latency during the day in both sexes (p<0.05; PA Figure 3I-J; Table S2). There were no significant effects of spermidine on sleep consolidation (p>0.05; Table S2). Spermidine did not increase attrition of either sex, and there were no significant effects of spermidine on activity index, suggesting that spermidine’s modest sleep effects are *bona fide* (p>0.05; Table S2).

Surprisingly, ornithine had no significant effect on any of the sleep or activity metrics tested in either sex, despite being tested at a dose well within the range previously reported to promote *Drosophila* sleep (p>0.05; Figure 3G-J; Table S2) ^1^. Our results suggest that ingested polyamines, but not ornithine, are somnogens in *Drosophila melanogaster*. While this is initially counter-intuitive given our RNAi results, we believe that homeostatic mechanisms that tightly regulate cellular polyamine levels may explain this seeming disconnect (see Discussion).

We next attempted to further our understanding of how polyamines regulate sleep by testing whether they could rescue sleep in our sleep mutants when supplemented. However, this proved impossible for an intriguing reason—sleep mutants were rapidly killed by introduction of polyamine-supplemented food over the course of our attempted sleep experiments. This prompted us to conduct an acute 6-day survival study of sleep mutant and control flies under these conditions, comparing within-genotype toxicity on ornithine, putrescine, and spermidine supplemented food vs vehicle control.

In females, spermidine was selectively toxic to *fmn* and *sss* mutants (p<0.0167; Figure4A-D). Females of all three sleep mutants also appeared to have elevated attrition rates on putrescine relative to *iso31* (Figure 4A-D). In males, putrescine was toxic to all three sleep mutants but not *iso31*, while spermidine was toxic only to *sss* flies (p<0.0001; Figure 4E-H). No sex-by-genotype combination showed toxicity over vehicle levels on ornithine (Figure 4A-H).

**Figure 4:**
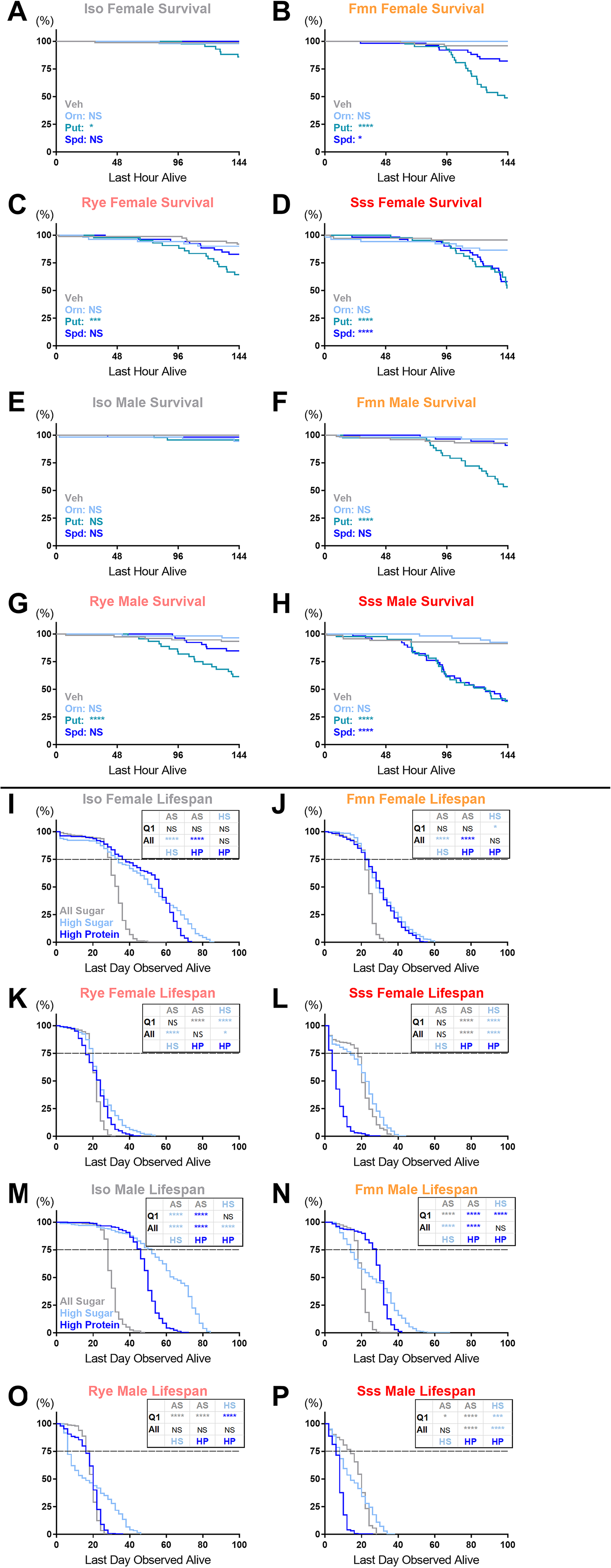
Dietary nitrogen is disproportionately toxic to *Drosophila* sleep mutants. (A-H) Acute survival of female (A-D) and male (E-H) flies during exposure to water vehicle (gray), 16mM ornithine (light blue), 16mM putrescine (medium blue), or 16mM spermidine (dark blue) supplemented sugar-agar in locomotor tubes. Genotypes are *iso31* (A,E), *fmn* (B,F), rye (C,G), and sss (D,H). n=41-72 flies per group; panels show percentage survival over time of whole population, censoring flies surviving >6 days; compared using within-genotype Wilcoxon tests of flies on each supplement vs vehicle. (I-P) Lifespan of female (I-L) and male (M-P) flies maintained on high-protein, high-sugar, and all-sugar diets. Genotypes are *iso31* (I,M), *fmn* (J,N), rye (K,O), and sss (L,P). n=206-239 flies per group; panels show lifespan as percentage of total population, excluding flies that escaped during vial flips; compared using within-genotype Wilcoxon tests of flies maintained on all 3 diets to each other. Comparisons were made assessing both acute toxicity of the diets (Q1: lifespan of shortest-lived quartiles in each group, censoring remaining three-quarters of the population; Q1 subpopulation is portion of each graph above the dashed line) and lifespan of the entire population on the diets (All: lifespan of the entire population considered). For all panels, significance threshold is adjusted to p=0.0167 after post-hoc Bonferroni correction for triplets of comparisons: NS = not significant, *p<0.0167, **p<0.0033; ***p<0.0003, ****p<0.0001; NS = not significant.

### Dietary protein disproportionately shortens the lifespan of *Drosophila* sleep mutants

The toxicity of polyamines in sleep mutants is consistent with sleep mutant sensitivity to dietary nitrogen. Indeed, according to previous reports *fmn* flies have normal lifespan on standard food, but shortened lifespan on a high-calorie diet with markedly higher percentage of calories from nitrogen-rich protein ^7,30^. However, shortened lifespan has previously been reported on standard food for *sss* flies, and is generally associated with chronic short-sleep ^2,9^. This raised the possibility that general ill-health might explain sleep mutant sensitivity to polyamines, rather than a specific effect stemming from altered nitrogen metabolism. Conducting a classical lifespan study with these mutants, we found that both females and males of all three genotypes (including *fmn*) have shortened lifespan on *iso31* background on standard fly food (p<0.0001; Figure S4A-B) ^7^. While this does not preclude sleep mutant sensitivity to nitrogen, we concluded that a paradigm allowing us to compare toxicity of nitrogen with that of another stressor was necessary to definitively determine whether sleep mutants are selectively vulnerable to nitrogen stress.

To accomplish this, we designed a lifespan study comparing within-genotype *iso31* and sleep mutant survival on calorie-matched: (1) high-protein food, (2) high-sugar food, and (3) all-sugar agar. A mostly-sugar diet with a small amount of protein is optimal for wild-type *Drosophila melanogaster* lifespan; thus, high-protein and all-sugar diets are both toxic to control flies compared to high-sugar diet ^31–35^. Based on our findings of disrupted nitrogen metabolism in sleep mutants, we expected their lifespan to be sensitized to high-protein and tolerant of all-sugar diet, relative to controls. We assessed both more acute toxicity in terms of death of the first quarter of each population (Q1 statistics) and chronic toxicity in the entire population (All statistics). We also conducted acute feeding assays with blue-dyed versions of these same diets, to assess whether differences in calorie intake might contribute to any lifespan effects we observed.

*iso31* females unexpectedly showed equivalent survival on high protein and high-sugar diets at the doses we used, although all-sugar was toxic (p<0.0001; Figure 4I). In contrast, sleep mutant females displayed both acutely (all mutants) and overall (*rye/sss*) accelerated death on high-protein compared to high-sugar diet (Figure 4J-L). Furthermore, all-sugar diet actually extended lifespan to varying degrees in *rye* and *sss* females (Figure 4K-L). Female flies fed similarly on all diets except for *sss*, which ate more high-sugar than high-protein food (Figure S4C-F). Thus, all diet effects on lifespan in *iso31, fmn*, and *rye* flies, and the protective effects of all-sugar in *sss*, lack a possible feeding confound.

Male *iso31* lifespan was longest on high-sugar diet, and longer on high-protein than on all-sugar diet (p<0.0001; Figure 4M). Surprisingly, high-protein diet was acutely protective in *fmn* and *rye* males, but this effect was not sustained with prolonged exposure (Figure 4N-O). However, all-sugar diet acutely extended lifespan vs high sugar (all three mutants) and high protein (*rye/sss*), and for both *rye* and *sss* the all-sugar diet was not the most toxic diet as it was for *iso31* controls (Figure 4M-P). The only feeding differences in males were in *iso31* and *sss* flies, which both ate less all-sugar compared to the other diets (Figure S4G-J). However, it is unlikely that calorie intake drives all-sugar longevity differences in *iso31* and *sss* males, as all-sugar diet has the same feeding effect and polar opposite lifespan effects in males of these genotypes (Figure 4M,P). The effects of all-sugar on *iso31* and *sss* males are also broadly similar to females, which lacked an all-sugar feeding confound.

Consistent with prior work on wild-type flies, diet composition rather than calorie intake appears to drive most, if not all, lifespan effects of diet in *iso31* and sleep mutant flies (Figures 4,S4) ^31–35^. Interestingly, longevity effects of diet composition are somewhat sexually dimorphic. While female sleep mutants are dose-dependently nitrogen sensitive, male sleep mutants require a more complete withdrawal of dietary nitrogen to promote survival, and *fmn* and *rye* males even appear to benefit in the short term from high-protein (Figure 4). That said, our nitrogen-enriched high-protein diet is toxic to varying degrees in all female sleep mutants but not *iso31* controls, while completely nitrogen-deficient all-sugar diet is protective to varying degrees in sleep mutants of both sexes, while profoundly shortening the lifespan of control flies (Figure 4). Overall, these results are consistent with toxicity of dietary nitrogen in chronically sleep deprived flies, similar to our observations with dietary polyamine supplementation.

## Discussion

Unbiased metabolomics of *Drosophila melanogaster* heads revealed two prominent metabolite clusters commonly dysregulated by chronic sleep loss across sleep mutants: (1) up-regulated ornithine, polyamines, and related stress-linked nitrogen metabolites, and (2) loss of lipids (especially cholesteryl esters, and many individual triacylglycerols), accumulated long-chain acyl-carnitines, and dysregulated TCA cycle metabolites, all suggestive of mitochondrial stress (Figures 1,S1; Table S1). Since no nitrogen pathway was consistently down-regulated at whose expense polyamines might be up-regulated (Table S1), we tested whether surfeit waste nitrogen could drive sleep mutant ornithine and polyamine elevation. Nitrogen inputs and waste products were not consistently elevated in all sleep mutants, but were consistently excreted much less efficiently than in controls, leading us to propose that elevated ornithine and polyamines in fly heads function as a nitrogen sink (Figures 2,S2). Both supplementing and blocking the synthesis of polyamines is sleep-promoting, suggesting that polyamine metabolism plays a complex role in sleep homeostasis (Figures 3). Finally, both polyamines and dietary protein are disproportionately toxic in sleep mutants, suggesting that the changes to their nitrogen metabolome contribute to their shortened lifespans (Figures 4). These findings are overlaid on a curated map of *Drosophila* nitrogen metabolic pathways in Figure 5.

**Figure 5:**
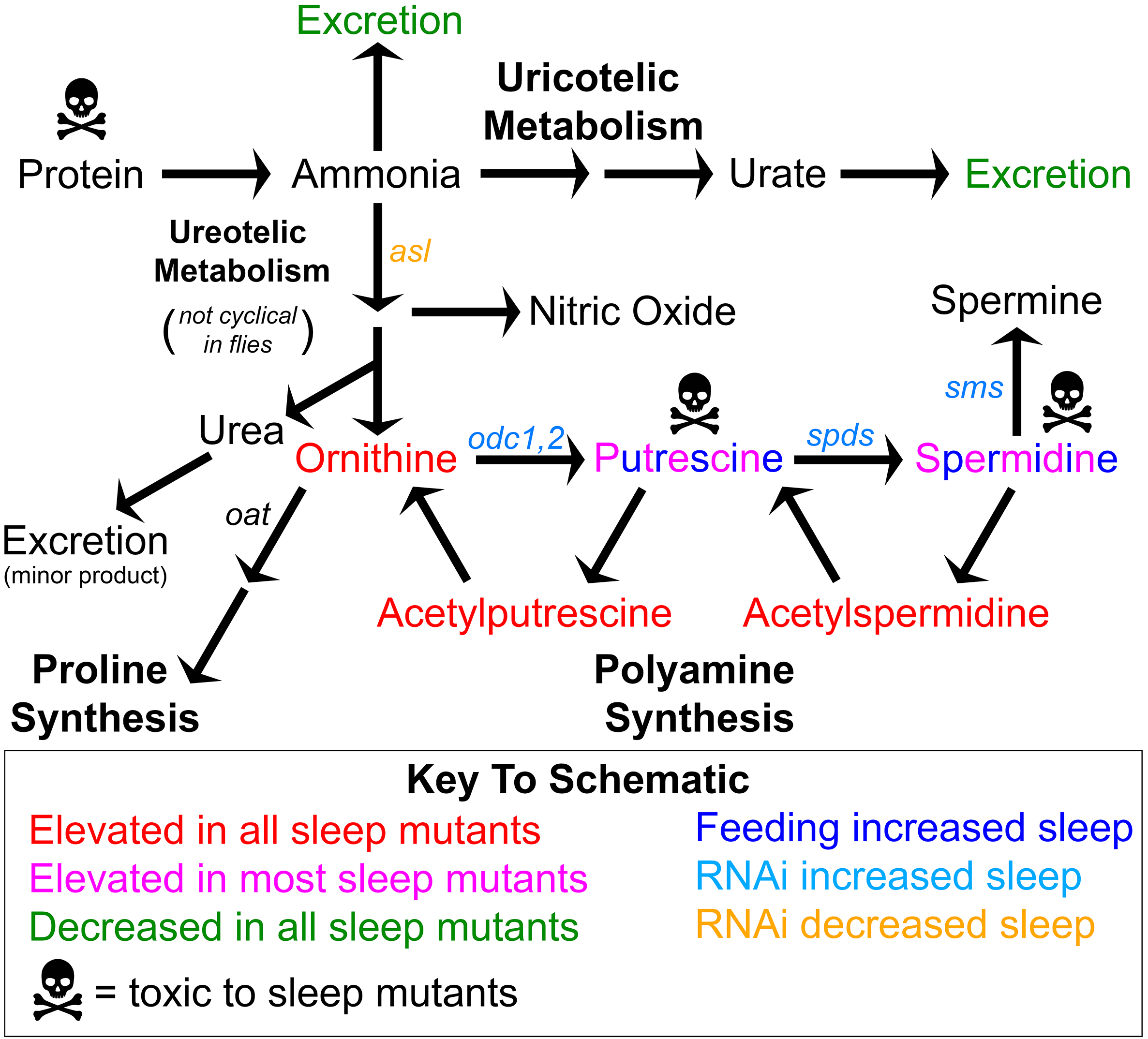
Schematic.

In this study, we sought to identify novel metabolic changes specific to chronic sleep loss as candidate effectors of sleep on lifelong health. Lipid and mitochondrial dysregulation very likely contribute to ill health in sleep mutants, given recent studies showing a central role of redox in linking sleep loss to lifespan ^36,37^. However, similar metabolic changes have been widely reported in both acute and chronic SD models. Acute SD modifies both expression of lipid-metabolism genes and levels of triacylglycerol, cholesterol, and/or acyl-carnitine species, often similarly to what we report here for our sleep mutants, in *Drosophila* ^5,38,39^, rodents ^5,40,41^, humans ^5,41–44^, and even “sleep-deprived” neurons in culture ^45^. Elevations in methylcitrate and decreased aconitate have also been reported, as have effects of lipid metabolism on sleep in *Drosophila* ^39,46–48^. Thus, we focused our follow-up studies on our other major finding.

To our knowledge, this is the first report linking chronic sleep loss to polyamine elevation in the tissues of any organism ^5^. However, there is indirect evidence that this extends to severe chronic sleep loss in humans. Sleep apnea patients excrete elevated levels of ornithine, putrescine, acetyl-ornithine, and acetyl-putrescine, as well as spermine in severe cases ^49,50^. Surveying numerous metabolomic studies in human and rodent models, we found no similar reports of polyamine elevation with more acute SD paradigms ^5^. Interestingly, in rat brain dialysate, acute SD alone has no effect on polyamines, but acute SD becomes able to increase putrescine in hyperammonemic rats ^51^. This supports the causal relationship we posit between deficient nitrogen waste clearance and polyamine accumulation in Drosophila sleep mutant heads (Figures 1,2) and its translatability to mammals. It is also worth noting that the selectivity of ornithine and polyamine, but not proline, metabolite up-regulation in chronic SD may be related to mitochondrial dysfunction implied by our other metabolomic findings. The mitochondrial enzyme *oat* gates entry of ornithine into proline synthesis, while the cytosolic/nuclear enzyme *odc* gates ornithine’s entry into polyamine synthesis ^52^. Mitochondrial dysfunction may thus favor the entry of excess ornithine into polyamine synthesis (Figure 5).

More isolated changes in ornithine level may be a leading indicator of nitrogen metabolome remodeling that is susceptible to shorter-term sleep loss than polyamines, though ornithine’s relationship to sleep is complex and sometimes contradictory. For example, ornithine is elevated in human plasma during SD according to one study, but during recovery sleep but not SD according to another ^53,54^. Our data are consistent with the latter pattern; ornithine alone among related nitrogen metabolites weakly trends higher during recovery sleep but not acute SD in *iso31* heads (Figure S1).

We also find that feeding polyamines, but not ornithine, increases sleep in *Drosophila melanogaster* (Figures 3G-J). Thus, long-term polyamine buildup from chronic SD may serve a sleep homeostatic function. However, it is important to note that we also observe sleep gain with RNAi knockdown of polyamine synthesis genes in the whole fly (*spds, sms*) and pan-neuronally (*odc1*) (Figures 3C-F). Since supplementing polyamines inhibits further production of polyamines from ornithine by increased antizyme translation and other well-conserved mechanisms, it is possible that feedback of polyamines onto *de novo* polyamine synthesis has somnogenic properties^24^ (Figure 5). Alternatively, deviations of polyamines outside of their usually-narrow physiological range may disrupt metabolic homeostasis, and sleep is required to restore the balance ^24^. In either case, our data implicate sleep homeostasis as a behavioral complement to the biochemical mechanisms that tightly control polyamine homeostasis at the cellular level. This is particularly intriguing given the relevance of polyamines to proposed functions of sleep, including roles in DNA repair, antioxidant properties, and regulation of macroautophagy ^2,13,24,36,37^.

Our finding that polyamines, but not ornithine, promote sleep is at odds with a prior report of ornithine supplementation promoting sleep selectively in mated female Drosophila ^1^. Genetic background effects could explain this discrepancy; for example, their fly strain may convert excess ornithine into sleep-promoting species at higher rates than *iso31*. However, we suspect they detected a confound from increased egg-laying and related chemotaxis, rather than sleep. Effects on egglaying have been directly shown for polyamines in *Drosophila*, and the polyamine precursors arginine and ornithine have similar reproduction-promoting effects in species where their roles have been examined ^29,55^. Use of position-agnostic multibeam monitors for our supplementation studies largely obviates this concern for our data, as do our findings of both female and male sleep gain on polyamines (Figure 3G-J).

Finally, we report a novel interaction between chronic sleep loss and diet that regulates fruit fly longevity (Figure 4). The toxicity of diets that deviate from the low protein-high carbohydrate optimum is well established for *Drosophila* of both sexes ^31–35^. We show that chronic sleep loss enhances protein toxicity in females, and actually changes an all-sugar diet from a toxic to a protective food substrate in both sexes (Figure 4). Importantly, the extent to which both polyamine feeding and dietary protein are toxic, and all-sugar diet is protective, in sleep mutants correlates with their degree of nitrogen metabolome remodeling. *fmn* flies exhibit milder polyamine elevation and nitrogen toxicity, while the phenotypes of *rye* and *sss* flies for all these metrics are more pronounced (Figures 1,4,S1). The toxic effect of high protein diets may explain why sleep-deprived humans crave food rich in carbohydrates; this could be an adaptive mechanism driving them towards safer food versus those that impair physiology under these conditions.

Upregulation of polyamines is particularly interesting in the context of recent work demonstrating that buildup of reactive oxygen species (ROS) in the fruit fly, especially in the gut, cause the shortened lifespan associated with chronic sleep loss ^36,37^. Polyamines themselves are antioxidants, and can further influence redox homeostasis by modulating substrate availability for producing urate, nitric oxide, and other relevant species ^24,56^. Moreover, deficits in nitrogen waste excretion in sleep mutants (Figure 2) may increase systemic oxidative stress ^57^. A promising direction for future research is whether nitrogen metabolism changes associated with chronic sleep loss interact with ROS buildup, particularly in the brain and gut ^36,37^.

Disordered nitrogen metabolism also has implications for aging-associated health outcomes beyond lifespan. Chronic SD is a well-known risk factor for neurodegeneration, but the mechanisms underlying this link are murky. Failure to excrete nitrogen during chronic SD likely predisposes the fly to systemic nitrogen stresses known to precede brain ROS accumulation and neurodegeneration in animal models ^58,59^. Furthermore, polyamine elevation in the head may play a local role in coupling chronic SD to neurodegeneration, although the relationship of polyamines to neurodegeneration is complex, as both protective ^60–63^ and pro-neurodegenerative ^64,65^ functions have been described. That said, relatively weaker induction of spermidine compared to more toxic putrescine and acetylated ‘back-converting’ polyamines by chronic SD hints that the overall effect may be more pro-than anti-neurodegenerative (Figures 1, S1).

Importantly, in the brain polyamines act in part through positive regulation of autophagy ^60,66^. Joint polyamine-autophagy mechanisms also play roles in several other aspects of cellular and organismal health ^67–69^. These connections are of particular interest to us, as we recently described a relationship between sleep and autophagy that may become maladaptive under chronic SD ^13^. It is tempting to speculate that remodeling of the nitrogen metabolome plays a role in this change. Both separately and together with autophagy, interactions between sleep and polyamines may contribute to pathology under conditions of long-term strain to polyamine homeostasis, such as aging; insomnia and other disorders causing chronic SD; and sustained metabolic stressors such as obesity and diabetes. Polyamine elevation and nitrogen retention, like autophagy, are also promising candidate mechanisms for linking sleep and food homeostasis over long timescales,

## Acknowledgements

We would like to thank Pavel Pivarshev for assistance with fly husbandry and handling.

## Figure Legends

**Figure S1:**
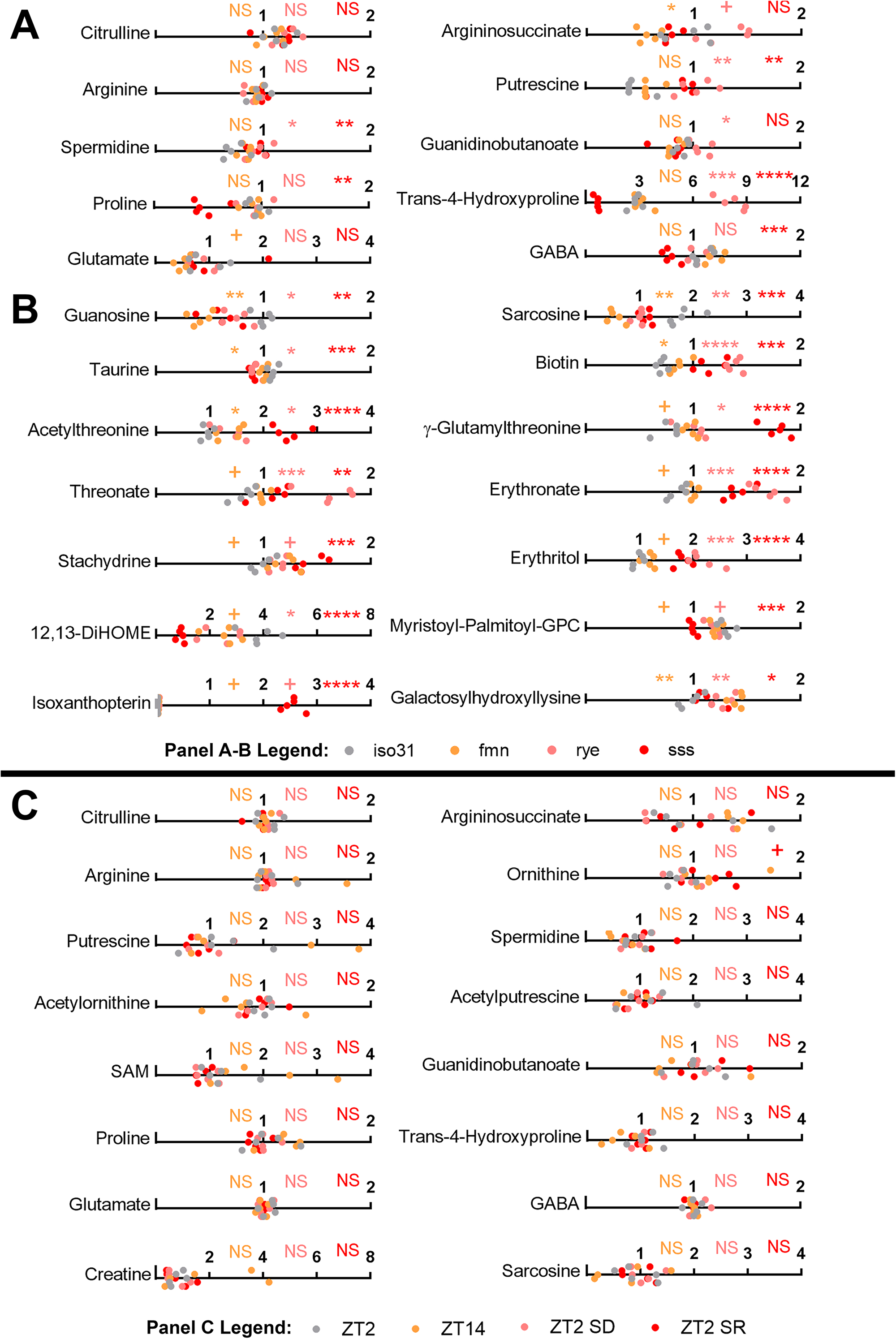
Additional metabolomic effects of chronic and acute sleep loss in *Drosophila*. All flies were maintained on a 12hr:12hr light:dark cycle during these experiments. (A-B) Line graphs showing scaled metabolite levels in *iso31* control (gray), *fmn* (orange), *rye* (pink), or *sss* (red) head lysates, grouped by involvement in nitrogen metabolism but with inconsistent effects across sleep mutants (A) or additional metabolites in assorted pathways that were consistently up- or down-regulated across sleep mutants (B). (C) Line graphs showing linear corrected metabolite levels in *iso31* control head lysates collected: at ZT2 (gray), at ZT14 (orange), after a ZT12-ZT2 mechanical sleep deprivation at ZT2 (pink), or after a ZT12-ZT0 sleep deprivation followed by ZT0-2 recovery sleep at ZT2 (red). This targeted metabolomics assay examined the levels of polyamines and metabolites in other pathways linked to polyamine synthesis. All data shown are individual pools of fly head lysate; n=5; Welch’s t-tests (p-values) with FDR correction for multiple comparisons (q-values); + p<0.05 but q>0.05, * p/q<0.05, ** p/q<0.01, *** p/q<0.001, **** p/q<0.0001.

**Figure S2:**
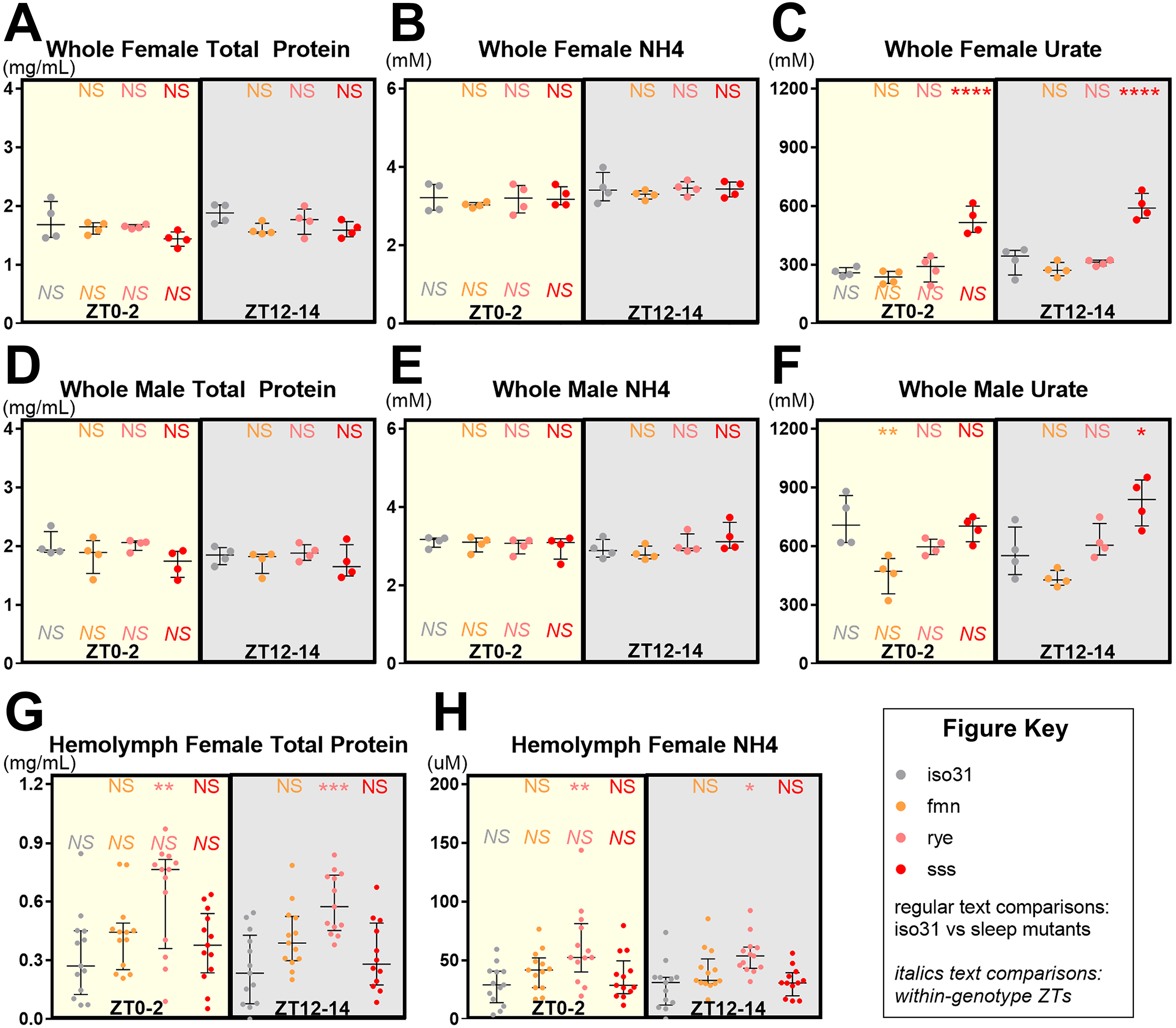
Sleep mutant total protein and waste nitrogen levels are not consistently altered. All biochemical data are from *iso31* (gray), *fmn* (orange), *rye* (pink), and *sss* (red) flies on a 12hr:12hr light:dark cycle. All top-row, regular-text statistical comparisons shown are for color-matched sleep mutant vs iso31 control. All bottom-row, italicized statistical comparisons shown are for color-matched within-genotype time-of-day comparisons. (A-F) Whole fly levels of total protein (A,D), ammonia (B,E), and urate (C,F) at ZT0-2 or ZT12-14 in female (A-C) and male (D-F) flies. Data shown are level in individual pools of whole fly lysate overlaid with median+/-interquartiles; n=4. (G-H) Hemolymph levels of total protein (G) and ammonia (H) at ZT0-2 or ZT12-14. Data shown are level in hemolymph bled from individual pools of female flies diluted in PBS + protease inhibitor, overlaid with median+/-interquartiles; n=12-13. All experiments used Tukey HSD tests; NS = not significant, * p<0.05, ** p<0.01, *** p<0.001, **** p<0.0001.

**Figure S3:**
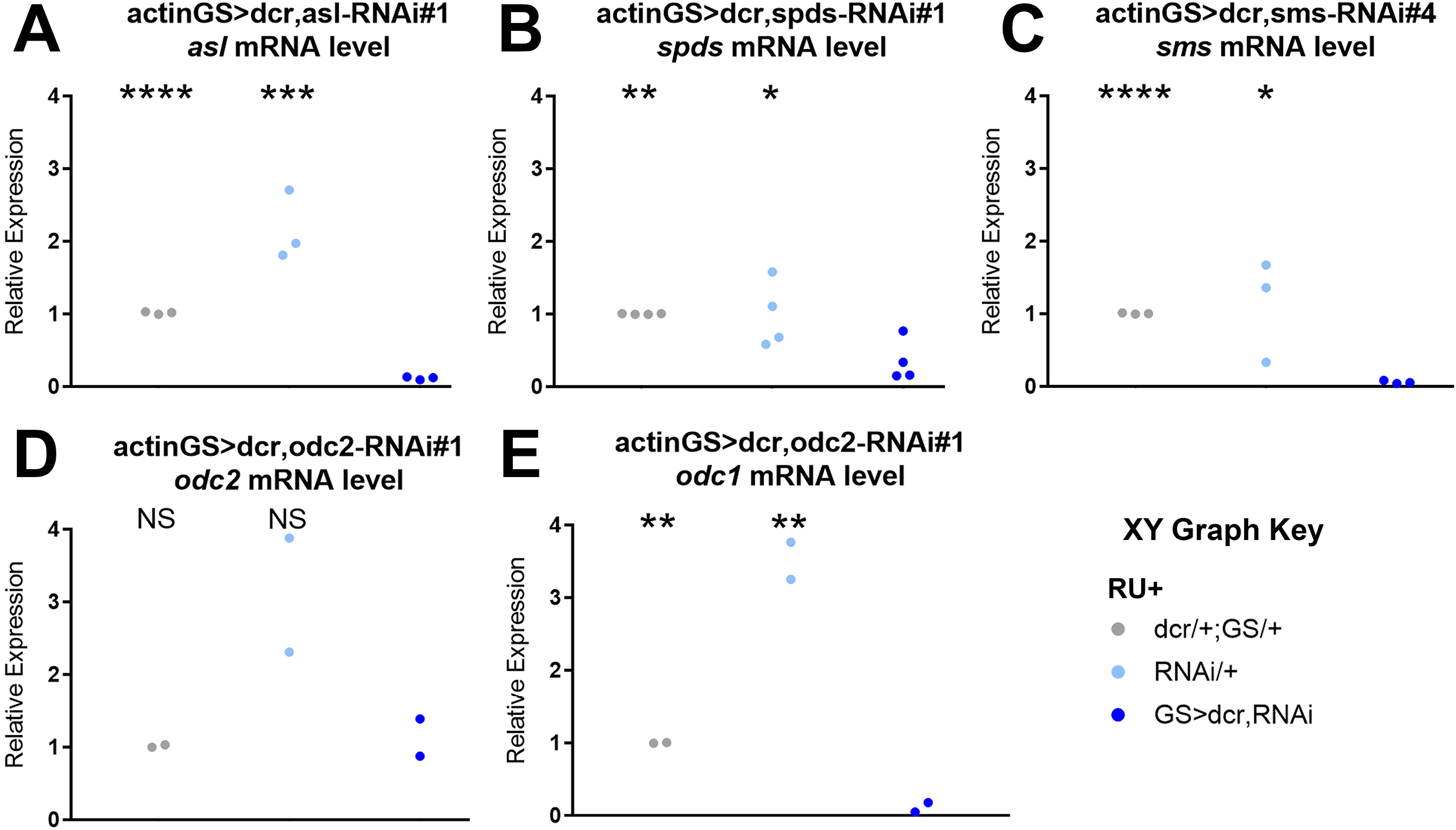
Efficacy of RNAi at knocking down target transcripts. (A-E) All panels show relative cDNA expression level measured by qPCR in whole-fly lysate of actinGS>dcr, RNAi flies compared to uas-dicer2/+;actinGS/+ and uas-RNAi/+ controls. We assess *asl* knockdown for asl-RNAi#1 (A), *spds* knockdown for spds-RNAi#1 (B), *sms* knockdown for sms-RNAi#4 (C) and both *odc2* and *odc1* knockdown for odc2-RNAi#1 (D-E). Data shown are relative expression level in individual pools of whole-fly lysate; n=2-4; one-tailed t-tests; all comparisons shown are for the corresponding control group vs actinGS>dcr,RNAi cross; NS = not significant, * p<0.05, ** p<0.01, *** p<0.001, **** p<0.0001.

**Figure S4:**
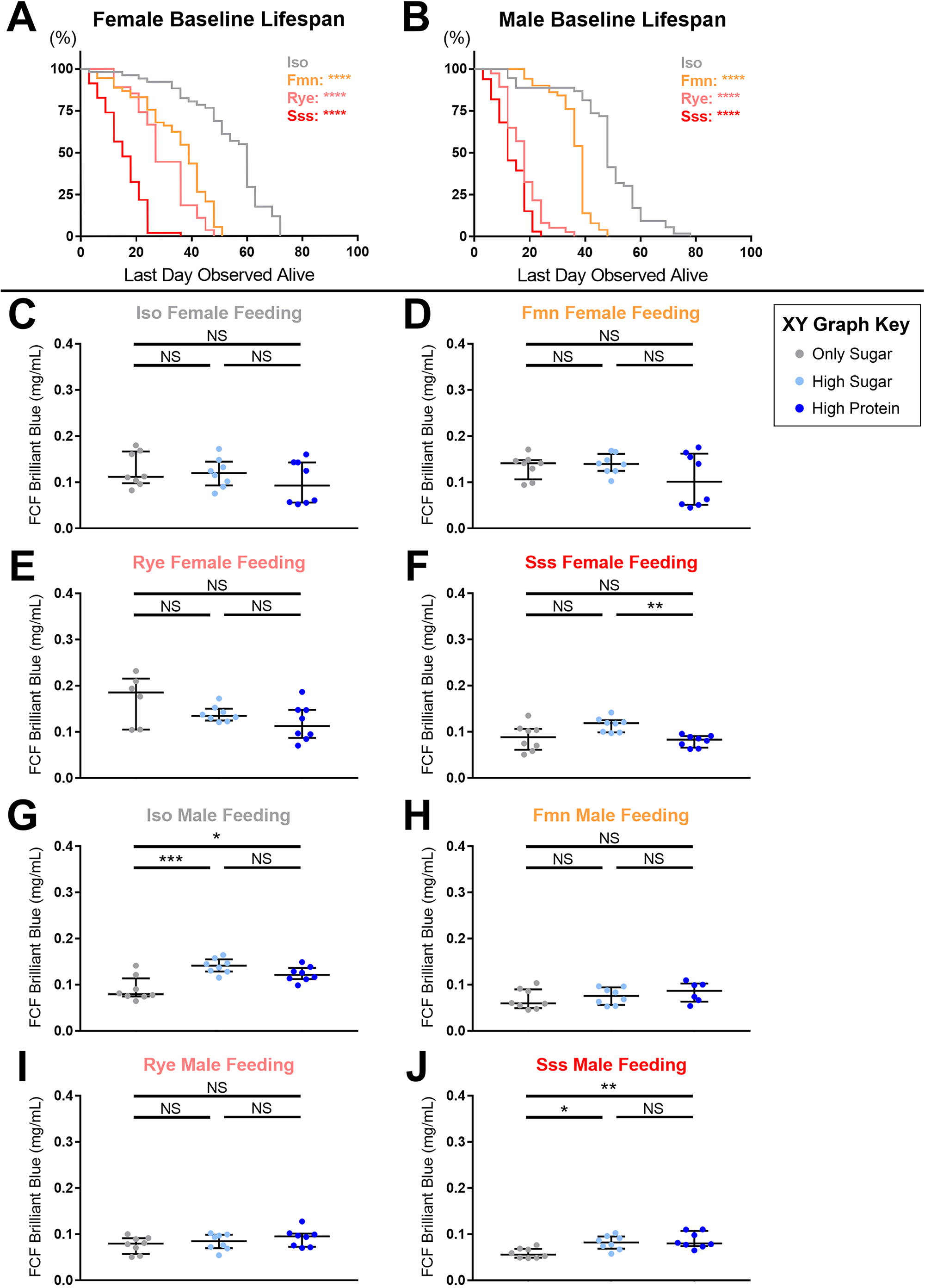
Baseline lifespan and feeding in sleep mutant and control flies. Flies were maintained on a 12hr:12hr light:dark cycle for the duration of these experiments. (A-B) Lifespan of female (A) and male (B) *iso31* (gray), *fmn* (orange), *rye* (pink), and *sss* (red) flies on our lab’s standard yeast-molasses food. Panels show lifespan as percentage of total population, excluding flies that escaped during vial flips; n=27-67 flies per group; Wilcoxon tests of each sleep mutant vs *iso31*. Significance threshold for (A-B) only is adjusted to p=0.0167 after post-hoc Bonferroni correction for triplets of comparisons: NS = not significant ; **** p<0.0001. (C-J) FCF brilliant blue content in whole fly lysates after 4hr acute feeding on high-protein, high-sugar, or all-sugar diet spiked with dye. Data from female (C-F) and male (G-J) flies of *iso31* control (C,G), *fmn* (D,H), *rye* (E,I), and *sss* (F,J) genotypes are shown. Panels show blue dye concentration in lysate collected from individual pools of flies overlaid with median+/- interquartiles; n=6-8; Tukey HSD tests; NS = not significant, * p<0.05, ** p<0.01, *** p<0.001.

<u>Table S1: Unbiased metabolomics screen of sleep mutant</u> *<u>Drosophila melanogaster</u>*.

**Tab#1**: sleep mutant / *iso31* control expression level of metabolites in assorted ureotelic and uricotelic nitrogen pathways, organized roughly in order within their respective pathways.

**Tab#2**: sleep mutant / *iso31* control expression levels of metabolites contributing to the Venn diagram in Figure 1A.

**Tab#3**: sleep mutant / iso31 control expression level of all metabolites from both the HD4 global metabolomics screen (top) and CLP lipidomics screen (bottom). Individual metabolites are loosely grouped by their predominant superpathway and subpathway groupings. p-values and q-values for each sleep mutant vs *iso31* control comparison can be found on the right.

**Tab#4**: scaled, imputed metabolite levels for each individual *iso31, fmn, rye*, and *sss* sample for both the HD4 (top) and CLP (bottom) screens.

**Tab#5**: original scale metabolite levels for each individual *iso31, fmn, rye*, and *sss* sample for both the HD4 (top) and CLP (bottom) screens.

<u>Table S2: Auxiliary sleep data from polyamine feeding and nitrogen metabolism RNAi sleep experiments.</u>

**Tab#1**: key listing stock center information and our own identifiers for each RNAi used for our sleep screens.

**Tab#2**: first-pass female total sleep mean, SEM, and n for each experimental and control group in our actinGS sleep screen. Difference of mean total sleep in each cross vs both control groups is also shown. Crosses that were considered first-pass hits (cross mean total sleep >=60min either higher or lower than both controls) are highlighted in gray. Crosses that survived validation and were identified as genuine hits are highlighted in red (sleep lower than controls) or blue (sleep higher than controls).

**Tab#3**: as Tab#4, but nsybGS screen.

**Tabs#4-7**: auxiliary sleep metrics including day/night sleep amount, activity index, mean bout length, and number of sleep bouts, as well as ZT0/ZT12 latency to sleep, are shown for each of the validated hits from our sleep screens. Median+/-interquartiles and n are shown for the experimental cross and each of the control groups, on both RU+ and RU-food (top). Statistical comparisons are shown for each control group compared to the experimental cross on each diet, and for within-genotype diet comparisons of each group (bottom).

**Tab#8**: female supplemental sleep metrics including day/night activity index, mean bout length, and number of sleep bouts, as well as ZT0/ZT12 latency to sleep. Median+/-interquartiles and n are shown for vehicle, ornithine, putrescine, and spermidine fed groups (top). Statistical comparisons are shown for all three nitrogen metabolites vs vehicle control (bottom).

**Tab#9**: as Tab#8, but with male flies.

## Literature Cited

1. Kanaya, H. J. et al. A sleep-like state in Hydra unravels conserved sleep mechanisms during the evolutionary development of the central nervous system. Sci. Adv. 6, eabb9415 (2020).

2. Mignot, E. Why We Sleep: The Temporal Organization of Recovery. PLoS Biol 6, e106 (2008).

3. Tononi, G. & Cirelli, C. Sleep function and synaptic homeostasis. Sleep Med Rev 10, 49–62 (2006).

4. Chouhan, N. S., Griffith, L. C., Haynes, P. & Sehgal, A. Availability of food determines the need for sleep in memory consolidation. Nature 589, 582–585 (2021).

5. Malik, D. M., Paschos, G. K., Sehgal, A. & Weljie, A. M. Circadian and Sleep Metabolomics Across Species. Journal of Molecular Biology 432, 3578–3610 (2020).

6. Dubowy, C. & Sehgal, A. Circadian Rhythms and Sleep in Drosophila melanogaster. Genetics 205, 1373–1397 (2017).

7. Kume, K., Kume, S., Park, S. K., Hirsh, J. & Jackson, F. R. Dopamine is a regulator of arousal in the fruit fly. J. Neurosci. 25, 7377–7384 (2005).

8. Shi, M., Yue, Z., Kuryatov, A., Lindstrom, J. M. & Sehgal, A. Identification of Redeye, a new sleep-regulating protein whose expression is modulated by sleep amount. Elife 3, e01473 (2014).

9. Koh, K. et al. Identification of SLEEPLESS, a sleep-promoting factor. Science 321, 372–376 (2008).

10. Chen, W.-F. et al. A neuron-glia interaction involving GABA transaminase contributes to sleep loss in sleepless mutants. Mol. Psychiatry 20, 240–251 (2015).

11. Murillo-Rodriguez, E. et al. Basic sleep mechanisms: an integrative review. Cent Nerv Syst Agents Med Chem 12, 38–54 (2012).

12. Seugnet, L. et al. Identifying sleep regulatory genes using a Drosophila model of insomnia. J Neurosci 29, 7148–7157 (2009).

13. Bedont, J. L. et al. Short and long sleeping mutants reveal links between sleep and macroautophagy. eLife 10, e64140 (2021).

14. Nieman, D. C., Gillitt, N. D., Sha, W., Esposito, D. & Ramamoorthy, S. Metabolic recovery from heavy exertion following banana compared to sugar beverage or water only ingestion: A randomized, crossover trial. PLoS ONE 13, e0194843 (2018).

15. Kuang, A. et al. Lipidomic Response to Coffee Consumption. Nutrients 10, E1851 (2018).

16. Ubhi, B. K. Direct Infusion-Tandem Mass Spectrometry (DI-MS/MS) Analysis of Complex Lipids in Human Plasma and Serum Using the Lipidyzer™ Platform. in Clinical Metabolomics (ed. Giera, M.) vol. 1730 227–236 (Springer New York, 2018).

17. Lintonen, T. P. I. et al. Differential Mobility Spectrometry-Driven Shotgun Lipidomics. Anal. Chem. 86, 9662–9669 (2014).

18. Baker, P. R. S., Armando, A. M., Campbell, J. L., Quehenberger, O. & Dennis, E. A. Three-dimensional enhanced lipidomics analysis combining UPLC, differential ion mobility spectrometry, and mass spectrometric separation strategies. Journal of Lipid Research 55, 2432–2442 (2014).

19. Rhoades, S. D. & Weljie, A. M. Comprehensive Optimization of LC-MS Metabolomics Methods Using Design of Experiments (COLMeD). Metabolomics 12, 183 (2016).

20. Malik, D. M., Rhoades, S. & Weljie, A. Extraction and Analysis of Pan-metabolome Polar Metabolites by Ultra Performance Liquid Chromatography-Tandem Mass Spectrometry (UPLC-MS/MS). Bio Protoc 8, e2715 (2018).

21. Hsu, C. T., Choi, J. T. Y. & Sehgal, A. Manipulations of the olfactory circuit highlight the role of sensory stimulation in regulating sleep amount. Sleep zsaa265 (2020) doi:10.1093/sleep/zsaa265.

22. Weihrauch, D. & O’Donnell, M. J. Mechanisms of nitrogen excretion in insects. Current Opinion in Insect Science 47, 25–30 (2021).

23. De Carvalho, F. G. et al. Taurine: A Potential Ergogenic Aid for Preventing Muscle Damage and Protein Catabolism and Decreasing Oxidative Stress Produced by Endurance Exercise. Front Physiol 8, 710 (2017).

24. Miller-Fleming, L., Olin-Sandoval, V., Campbell, K. & Ralser, M. Remaining Mysteries of Molecular Biology: The Role of Polyamines in the Cell. Journal of Molecular Biology 427, 3389–3406 (2015).

25. Yan, L.-J., Levine, R. L. & Sohal, R. S. Oxidative damage during aging targets mitochondrial aconitase. Proceedings of the National Academy of Sciences 94, 11168–11172 (1997).

26. Fu, M.-X. et al. The Advanced Glycation End Product, N∈-(Carboxymethyl)lysine, Is a Product of both Lipid Peroxidation and Glycoxidation Reactions. Journal of Biological Chemistry 271, 9982–9986 (1996).

27. Amaral, A. U., Cecatto, C., Castilho, R. F. & Wajner, M. 2-Methylcitric acid impairs glutamate metabolism and induces permeability transition in brain mitochondria. J Neurochem 137, 62–75 (2016).

28. Melo, D. R., Mirandola, S. R., Assunção, N. A. & Castilho, R. F. Methylmalonate impairs mitochondrial respiration supported by NADH-linked substrates: Involvement of mitochondrial glutamate metabolism. J. Neurosci. Res. 90, 1190–1199 (2012).

29. Hussain, A. et al. Ionotropic Chemosensory Receptors Mediate the Taste and Smell of Polyamines. PLoS Biol 14, e1002454 (2016).

30. Yamazaki, M. et al. High calorie diet augments age-associats sleep impairment in Drosophila. Biochemical and Biophysical Research Communications 417, 812–816 (2012).

31. Jensen, K., McClure, C., Priest, N. K. & Hunt, J. Sex-specific effects of protein and carbohydrate intake on reproduction but not lifespan in Drosophila melanogaster. Aging Cell 14, 605–615 (2015).

32. Skorupa, D. A., Dervisefendic, A., Zwiener, J. & Pletcher, S. D. Dietary composition specifies consumption, obesity, and lifespan in Drosophila melanogaster. Aging Cell 7, 478–490 (2008).

33. Bruce, K. D. et al. High carbohydrate-low protein consumption maximizes Drosophila lifespan. Exp Gerontol 48, 1129–1135 (2013).

34. Lee, K. P. et al. Lifespan and reproduction in Drosophila: New insights from nutritional geometry. Proc Natl Acad Sci U S A 105, 2498–2503 (2008).

35. Mair, W., Piper, M. D. W. & Partridge, L. Calories do not explain extension of life span by dietary restriction in Drosophila. PLoS Biol 3, e223 (2005).

36. Hill, V. M. et al. A bidirectional relationship between sleep and oxidative stress in Drosophila. PLoS Biol. 16, e2005206 (2018).

37. Vaccaro, A. et al. Sleep Loss Can Cause Death through Accumulation of Reactive Oxygen Species in the Gut. Cell 181, 1307–1328.e15 (2020).

38. Cirelli, C., LaVaute, T. M. & Tononi, G. Sleep and wakefulness modulate gene expression in Drosophila. Journal of Neurochemistry 94, 1411–1419 (2005).

39. Thimgan, M. S., Seugnet, L., Turk, J. & Shaw, P. J. Identification of genes associated with resilience/vulnerability to sleep deprivation and starvation in Drosophila. Sleep 38, 801–814 (2015).

40. Cirelli, C., Gutierrez, C. M. & Tononi, G. Extensive and Divergent Effects of Sleep and Wakefulness on Brain Gene Expression. Neuron 41, 35–43 (2004).

41. Weljie, A. M. et al. Oxalic acid and diacylglycerol 36:3 are cross-species markers of sleep debt. Proc. Natl. Acad. Sci. U.S.A. 112, 2569–2574 (2015).

42. Davies, S. K. et al. Effect of sleep deprivation on the human metabolome. Proceedings of the National Academy of Sciences 111, 10761–10766 (2014).

43. Chua, E. C.-P., Shui, G., Cazenave-Gassiot, A., Wenk, M. R. & Gooley, J. J. Changes in Plasma Lipids during Exposure to Total Sleep Deprivation. Sleep 38, 1683–1691 (2015).

44. van den Berg, R. et al. A single night of sleep curtailment increases plasma acylcarnitines: Novel insights in the relationship between sleep and insulin resistance. Archives of Biochemistry and Biophysics 589, 145–151 (2016).

45. Hinard, V. et al. Key Electrophysiological, Molecular, and Metabolic Signatures of Sleep and Wakefulness Revealed in Primary Cortical Cultures. Journal of Neuroscience 32, 12506–12517 (2012).

46. Pamboro, E. L. S., Brown, E. B. & Keene, A. C. Dietary fatty acids promote sleep through a taste-independent mechanism. Genes, Brain and Behavior 19, (2020).

47. Thimgan, M. S., Suzuki, Y., Seugnet, L., Gottschalk, L. & Shaw, P. J. The perilipin homologue, lipid storage droplet 2, regulates sleep homeostasis and prevents learning impairments following sleep loss. PLoS Biol. 8, (2010).

48. Thimgan, M. S., Kress, N., Lisse, J., Fiebelman, C. & Hilderbrand, T. The acyl-CoA Synthetase, pudgy, Promotes Sleep and Is Required for the Homeostatic Response to Sleep Deprivation. Front. Endocrinol. 9, 464 (2018).

49. Xu, H. et al. Metabolomics Profiling for Obstructive Sleep Apnea and Simple Snorers. Sci Rep 6, 30958 (2016).

50. Xu, H. et al. Pediatric Obstructive Sleep Apnea is Associated With Changes in the Oral Microbiome and Urinary Metabolomics Profile: A Pilot Study. Journal of Clinical Sleep Medicine 14, 1559–1567 (2018).

51. Marini, S. et al. Abnormalities in the Polysomnographic, Adenosine and Metabolic Response to Sleep Deprivation in an Animal Model of Hyperammonemia. Front Physiol 8, 636 (2017).

52. Schipper, R. G. & Verhofstad, A. A. J. Distribution Patterns of Ornithine Decarboxylase in Cells and Tissues: Facts, Problems, and Postulates. J Histochem Cytochem. 50, 1143–1160 (2002).

53. Grant, L. K. et al. Circadian and wake-dependent changes in human plasma polar metabolites during prolonged wakefulness: A preliminary analysis. Sci Rep 9, 4428 (2019).

54. Honma, A. et al. Effect of acute total sleep deprivation on plasma melatonin, cortisol and metabolite rhythms in females. Eur J Neurosci 51, 366–378 (2020).

55. Lefèvre, P. L. C., Palin, M.-F. & Murphy, B. D. Polyamines on the reproductive landscape. Endocr Rev 32, 694–712 (2011).

56. Alvarez-Lario, B. & Macarron-Vicente, J. Uric acid and evolution. Rheumatology 49, 2010–2015 (2010).

57. Bosoi, C. R. et al. Systemic oxidative stress is implicated in the pathogenesis of brain edema in rats with chronic liver failure. Free Radical Biology and Medicine 52, 1228–1235 (2012).

58. Qvartskhava, N. et al. Hyperammonemia in gene-targeted mice lacking functional hepatic glutamine synthetase. Proc Natl Acad Sci U S A 112, 5521–5526 (2015).

59. Bergin, D. H. et al. Altered plasma arginine metabolome precedes behavioural and brain arginine metabolomic profile changes in the APPswe/PS1ΔE9 mouse model of Alzheimer’s disease. Transl Psychiatry 8, 108 (2018).

60. Gupta, V. K. et al. Restoring polyamines protects from age-induced memory impairment in an autophagy-dependent manner. Nat Neurosci 16, 1453–1460 (2013).

61. Büttner, S. et al. Spermidine protects against α-synuclein neurotoxicity. Cell Cycle 13, 3903–3908 (2014).

62. Phadwal, K. et al. Spermine increases acetylation of tubulins and facilitates autophagic degradation of prion aggregates. Sci Rep 8, 10004 (2018).

63. Luo, J. et al. Endogenous polyamines reduce the toxicity of soluble aβ peptide aggregates associated with Alzheimer’s disease. Biomacromolecules 15, 1985–1991 (2014).

64. Lewandowski, N. M. et al. Polyamine pathway contributes to the pathogenesis of Parkinson disease. Proc Natl Acad Sci U S A 107, 16970–16975 (2010).

65. Sandusky-Beltran, L. A. et al. Spermidine/spermine-N1-acetyltransferase ablation impacts tauopathy-induced polyamine stress response. Alzheimers Res Ther 11, 58 (2019).

66. Li, C. et al. Spermine synthase deficiency causes lysosomal dysfunction and oxidative stress in models of Snyder-Robinson syndrome. Nat Commun 8, 1257 (2017).

67. Eisenberg, T. et al. Induction of autophagy by spermidine promotes longevity. Nat. Cell Biol. 11, 1305–1314 (2009).

68. Zwighaft, Z. et al. Circadian Clock Control by Polyamine Levels through a Mechanism that Declines with Age. Cell Metabolism 22, 874–885 (2015).

69. Zhang, H. et al. Polyamines Control eIF5A Hypusination, TFEB Translation, and Autophagy to Reverse B Cell Senescence. Molecular Cell 76, 110–125.e9 (2019).

